# An optimal growth law for RNA composition and its partial implementation through ribosomal and tRNA gene locations in bacterial genomes

**DOI:** 10.1101/2021.02.05.429890

**Authors:** Xiao-Pan Hu, Martin J. Lercher

## Abstract

The distribution of cellular resources across bacterial proteins has been quantified through phenomenological growth laws. Here, we describe a complementary bacterial growth law for RNA composition, emerging from optimal cellular resource allocation into ribosomes and ternary complexes. The predicted decline of the tRNA/rRNA ratio with growth rate agrees quantitatively with experimental data. Its regulation appears to be implemented in part through chromosomal localization, as rRNA genes are typically closer to the origin of replication than tRNA genes and thus have increasingly higher gene dosage at faster growth. At the highest growth rates in *E. coli*, the tRNA/rRNA ratio appears to be regulated entirely through this effect.

## Background

The systematic change of the coarse-grained composition of bacterial proteomes with growth rate [1,2] can be quantified through phenomenological growth laws [3,4]. The most prominent growth law describes an apparently linear increase of the ribosomal protein fraction with growth rate [1,3]. These laws have been successfully applied to the prediction of a range of phenotypic observations [3,5–8]. Recently, it has been argued that they arise from an optimal balance between the cellular investment into catalytic proteins and their substrates [9].

In contrast to the proteome composition, the partitioning of bacterial RNA into messenger (mRNA), ribosomal (rRNA), and transfer (tRNA) RNA is often assumed to be growth rate-independent [2,3,5,6,10,11]. However, experimental evidence suggests that the tRNA/rRNA expression ratio decreases monotonically with growth rate [12–20], suggesting the existence of a bacterial growth law for RNA composition.

The regulatory implementation of bacterial growth laws is generally assumed to arise from a small number of major transcriptional regulators such as ppGpp [21,22] and cAMP [4,23]. However, growth-rate dependent transcriptional regulation could also be implemented through chromosomal gene positioning. Prokaryotic genes are non-randomly located on multiple levels [24–26], with highly expressed genes biased towards the origin of replication (oriC) [27]. The latter observation is thought to facilitate high expression levels at fast growth, where, due to the presence of partially replicated chromosomes, gene copy numbers depends on the distance to oriC (replication-associated gene dosage) [28–30]. Indeed, chromosome rearrangements that shift highly expressed genes from the origin to the terminus of replication reduce fitness [31–35].

rRNA forms the central part of the catalyst of peptide elongation, while tRNA forms the core of the substrate; together, they account for the bulk of cellular RNA [2]. Their cytosolic concentrations at different growth rates in *E. coli* are well described by an optimality assumption [9,36,37]. Moreover, chromosomal gene positions in *E. coli* are known to affect the expression of both tRNA and rRNA genes [38,39]; both types of genes are located closer to oriC in fast compared to slowly growing bacteria, with rRNA genes positioned closer to oriC than tRNA genes in most examined fast-growing bacteria [27].

Based on these previous observations, we hypothesize (i) that the relative expression of tRNA and rRNA can be described by a bacterial growth law that arises from optimal resource allocation and (ii) that this growth law is at least partially implemented through the relative chromosomal positioning of tRNA and rRNA genes.

## Results and discussion

In terms of dry mass allocation, translation is the most expensive process in fast-growing bacteria [2,40]. As evidenced by comparison of diverse data to a detailed biochemical model of translation, the allocation of cellular resources across components of the *E. coli* translation system minimizes their total dry mass concentration at a given protein production rate [37]. This result indicates that natural selection favored the parsimonious allocation of cellular resources to the translation machinery in *E. coli*. To generalize this optimization hypothesis to other species, we here analyze a coarse-grained translation model that only considers peptide elongation, where the active ribosome acts as an enzyme that converts ternary complexes (TC), consisting of elongation factor Tu (EF-Tu), GTP, and charged tRNA, into an elongating peptide chain following Michaelis-Menten kinetics [5,41]. In exponential, balanced growth at rate *μ* and cellular protein concentration *P*, the total rate of protein production is *v* = *μ · P*. We derived the optimal concentration ratio of TC (with molecular mass *m*_TC_) to ribosome (*R*, with molecular mass *m*_R_) at this production rate by minimizing their combined mass concentration, *M*_total =_*m*_TC_[TC] +*m*_R_[*R*] (**Methods**):

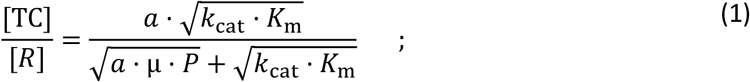

Here, 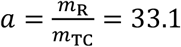 is the ratio of molecular weights of ribosome and TC; k_cat_ is the turnover number of the ribosome; and *K*_m_ is the ribosome’s Michaelis constant for TC, which is limited by diffusion [5]. For a given genome, *a, k*_cat_, and *K*_m_ can be treated as constants [5,6]. Moreover, the cellular protein mass concentration *P* appears to be similar across most species [42] and shows only minor variations across growth rates in those bacteria where it has been tested [7,43,44]. Thus, equation (1) predicts that the TC/ribosome ratio is a monotonically decreasing function of the growth rate *μ*. Since most cellular EF-Tu and tRNA are present in the form of TCs [5], hereafter, the TC concentration is assumed to be approximately equal to the concentration of EF-Tu and tRNA.

To calculate the optimal TC/ribosome ratio in *E. coli*, we use the measured protein concentration *P* [45], set the turnover number *k*_cat_ to the maximal observed translation rate [2], and set the Michaelis constant *K*_m_ to its diffusion limit [5] (Methods; see also Ref. [37]). **Fig. 1a** compares the optimal predictions (red line) to experimental datasets that estimated the TC/ribosome ratio based on ratios of tRNA/rRNA [20,46,47], EF-Tu/rRNA [19], and EF-Tu/ribosomal proteins [45] (**Additional file 1: Table S1**). Consistent with the predictions, all experimental estimates of the TC/ribosome ratio are approximately two-fold higher at low compared to high growth rates. As the TC and ribosome constitute the two major components of cellular RNA [2],we conclude that the optimal TC/ribosome ratio according to equation (1) represents a bacterial growth law for RNA composition:

**Figure 1.**
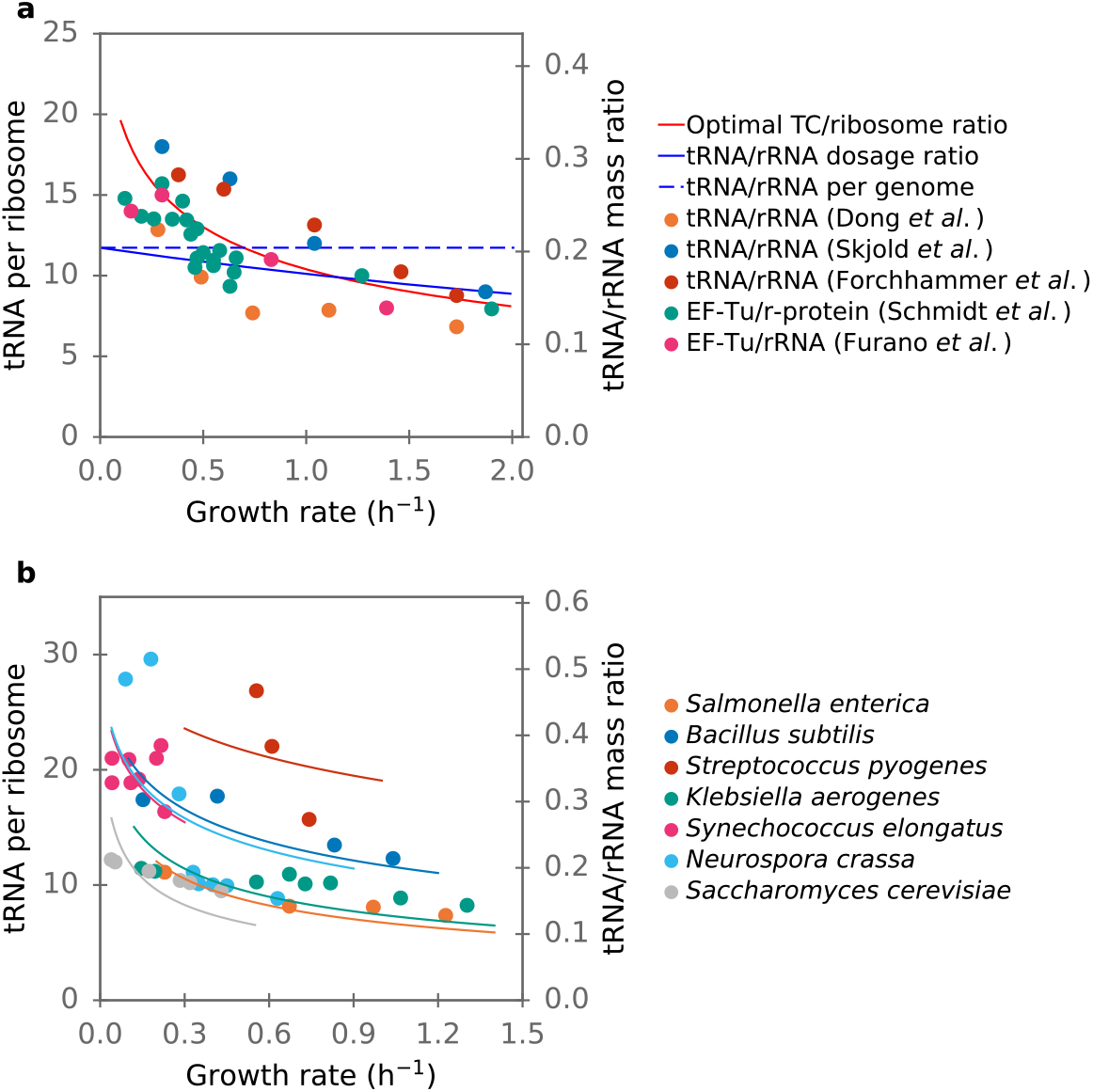
The RNA composition growth law predicts decreasing TC/ribosome ratios with growth rate. (**a**) Different experimental estimates of TC/ribosome ratios in *E. coli* are consistent with the optimal ratio according to equation (1) (red line). The solid blue curve marks the replication-associated tRNA/rRNA dosage ratio based on chromosomal positions of tRNA and rRNA genes; the dashed blue line marks the expectation based on the tRNA/ribosome gene ratio per chromosome. (**b**) Experimental estimates of TC/ribosome ratios in different microbes are also largely consistent with the optimal ratio; for each dataset, we fitted equation (1) to the data by varying the single adjustable parameter *k* (colored lines).

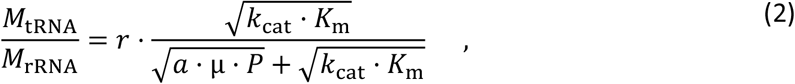

where *M*_tRNA_ and *M*_rRNA_ are the cellular mass of tRNA and rRNA, respectively, and *r*=0.58 is the ratio of the tRNA mass fraction of a TC and the rRNA mass fraction of the bacterial ribosome (**Methods**).

The approximate Michaelis-Menten form of the rate law for peptide elongation, on which the above equations are based, arises from the structure of the detailed elongation process [41]. As this process is shared by all living cells [41], we expect that the RNA composition growth law (2) holds also for other fast-growing microbes (with *a*=40.3 and *r*=0.59 in eukaryotes). To test this hypothesis, we collected all available tRNA/rRNA ratios in microbes (**Fig. 1b** and **Additional file 1: Table S2**). Note that if protein concentration *P* is indeed approximately constant across species [42], then equations (1) and (2) contain a single species-specific parameter, the geometric mean of *k*_cat_ and *K*_m_,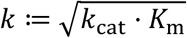.

For six out of the seven datasets in **Fig. 1**, the tRNA/ribosome ratio decreases with increasing growth rate. The only exception, the cyanobacterium *Synechococcus elongatus*, has a much smaller maximal growth rate (μ_max_=0.23h^-1^) than the other species, and its tRNA/rRNA ratio does not show a clear growth rate-dependence [48]. It is conceivable that slow-growing species do not fully optimize their translation machinery composition, as a near-optimal constant TC/ribosome ratio may incur a lower fitness cost than a regulatory system for growth rate-dependent expression.

We define gene dosage as the DNA copy number of a given gene in one cell. Gene dosage can change permanently because of chromosomal duplications and deletions, but also transiently according to the progress of chromosome replication. The latter causes a gene position-dependent dosage effect, with genes closer to the origin of replication (oriC) having a higher average dosage than genes further away (equation (14)). Gene dosage effects are in particular known to affect the expression of rRNA [38] and tRNA genes [39]. Moreover, a strong selection pressure towards optimal tRNA/ribosome ratios appears to exist in fast-growing bacteria (**Fig 1b**). We thus hypothesized that the relative genomic position of tRNA and rRNA genes may contribute to the implementation of the RNA composition growth law (**Fig 2a**). Here and below, for each genome, we summarize the multiple tRNA genes by averaging over their positions; we do the same for the rRNA genes.

**Figure 2.**
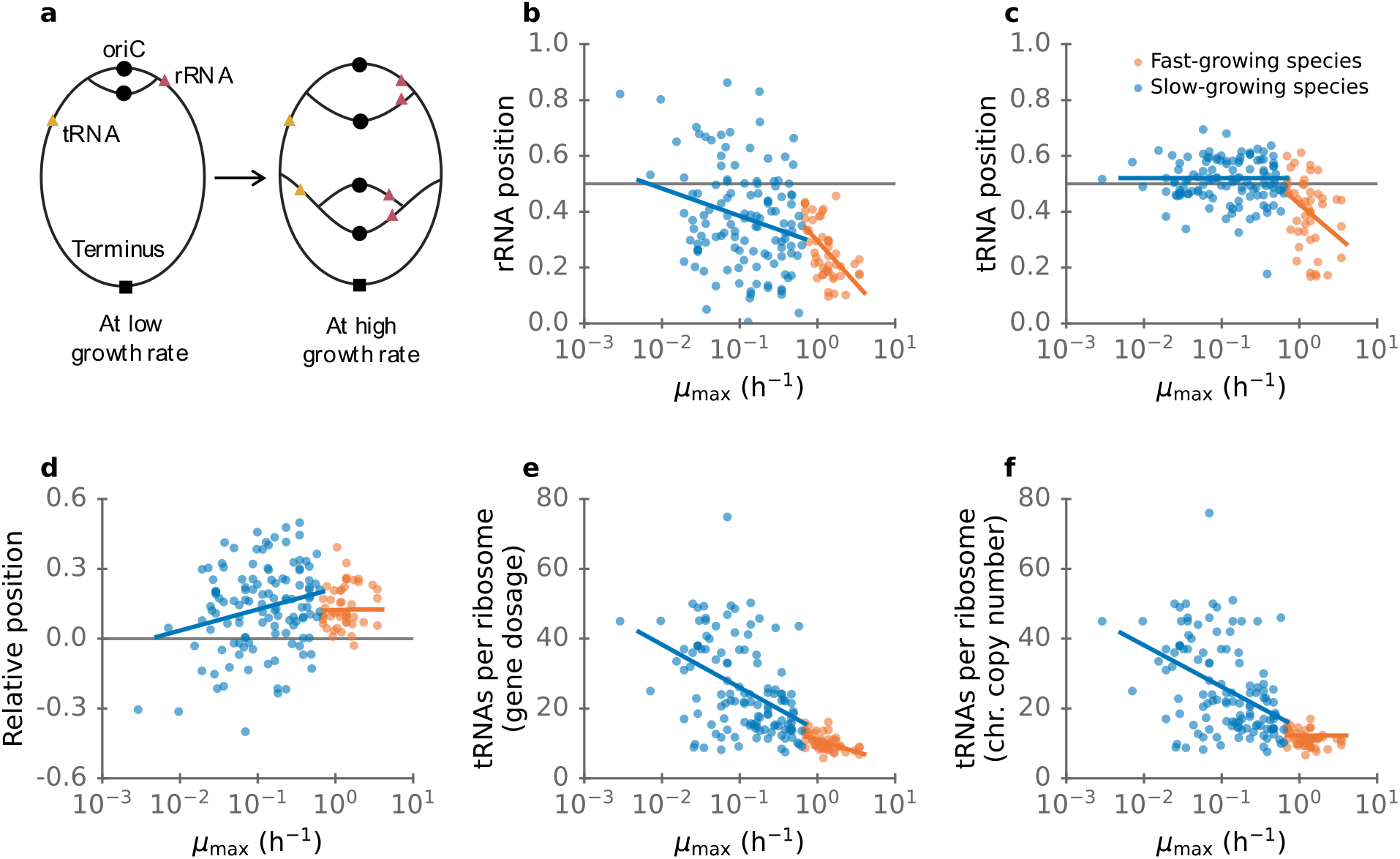
The genomic positioning of tRNA and rRNA genes contributes to the regulation of RNA composition. **(a)** A schematic diagram showing the dosage ratio of two genes as a function of growth rate, quantified in equations (3) and (16). (**b**) Arithmetic means of the tRNA positions for individual genomes as a function of *μ_max_*. The horizontal grey line marks the midpoint between origin and terminus of replication. (**c**) Same for rRNA. (**d**) Relative positions between tRNA and rRNA genes (position_tRNA_-position_rRNA_). The horizontal grey line marks the relative position 0. (**e**) tRNA/rRNA dosage ratios. (**f**) Genomic tRNA/rRNA ratios. In (b)-(f), blue points indicate slow growing species (with blue linear regression line) and orange points indicate fast-growing species (with orange linear regression line).

In line with previous analyses [27,49], we found that in fast-growing species (minimal doubling time ≤ 1 h, i.e., *μ*_max._ ≥0.69 h^−1^), rRNA and tRNA genes are generally located in the vicinity of oriC, at relative positions <0.5 (**Fig. 2b,c**, orange points; *μ*_max_ from Ref. [49]). Moreover, we found that both rRNA and tRNA genes tend to be located ever closer to oriC with increasing *μ*_max_ (Spearman’s rank correlation coefficient between *μ*_max_ and position_rRNA_: *ρ* = −0.59, *P* = 9.2 × 10^−6^; between *μ*_max_ and position_tRNA_: *ρ* = −=0.40, *P* = 0.0047). In slow-growing species, rRNA genes still tend to be close to oriC (**Fig. 2b**, blue points), while tRNA genes are distributed around the midpoint between oriC and the terminus (**Fig. 2c**, blue points). We found that rRNA genes are closer to oriC than tRNA genes in most slow-growing and in all but one fast-growing bacteria (**Fig. 2d**). To analyze how this affects the tRNA/rRNA ratio, we used the model developed by Bremer and Churchward [30] for the dosage ratio of two genes at growth rate *μ*,

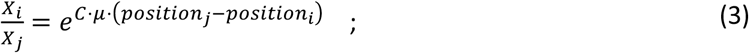

here, *X_i_*is the dosage and *position_i_*is the position of gene *i*, and C is the time required to complete one round of chromosome replication (see Methods for details). The DNA replication rate varies between species but is approximately constant across conditions in a given species [50], making the gene dosage ratio a monotonous function of the growth rate [51,52]. For tRNA and rRNA with multiple genes, the overall dosage ratio can be described by a generalization of equation (3) (Methods, equation (16)). As rRNA gene positions tend to be smaller than tRNA positions (**Fig. 2d**), the tRNA/rRNA ratio that would be obtained if regulation was exclusively by gene dosage is a decreasing function of growth rate, in qualitative agreement with the optimality predictions from equation (2).

This finding supports our hypothesis that natural selection has fine-tuned the positions of tRNA and rRNA genes to match the RNA composition growth law for optimally efficient translation. This can best be seen for *E. coli*, for which all necessary parameters are available, allowing us to make quantitative predictions without adjustable parameters (**Fig. 1a**). Here, the tRNA/rRNA dosage ratio at higher growth rates (1h^−1^ ≤ *μ* ≤ 2h^−1^) is very close to the optimal prediction, which corresponds to about 9 tRNAs per ribosome. This result is consistent with the notion that at the highest growth rates, both tRNA and rRNA genes are transcribed at the maximally possible rate, such that their relative expression is indeed dominated by gene dosage effects. The expression of both tRNA and rRNA operons is regulated by the P1 promoter, which is repressed by ppGpp; at near-maximal growth rates, ppGpp concentrations are low and the P1 promoter works near its maximal capacity [53]. In contrast, at low growth rates, P1 is repressed by ppGpp, and thus gene dosage can only partially explain the tRNA/rRNA ratio in these conditions.

According to equation (1), faster growing species need a lower TC/ribosome ratio at maximal growth. We indeed find statistically highly significant negative correlations between the tRNA/ribosome ratio and *μ_max_*(**Fig. 2e**; slowly growing species: *ρ* = −0.44, *P* = 2.8 × 10^−7^; fast-growing species: *ρ* = −0.49, *P* = 0.00043). While slowly growing species show a wide range of tRNA/ribosome gene dosage ratios, the ratio in fast-growing species shows a much tighter distribution. The chromosomal tRNA/ribosome copy number ratios (**Fig. 2f**) in slowly growing species are almost identical to the gene dosage ratios. Chromosomal gene copy number ratios in fast-growing species show a similarly tight distribution as gene dosage ratios (with 10^th^ percentile = 8.78 and 90^th^ percentile = 14.5), they show no strong systematic dependence on *μ_max_*(**Fig. 2e**, *ρ* = −0.24, *P* = 0.10). Interestingly, we also find no statistically significant dependence of the relative position on *μ_max_*in fast-growing species (**Fig. 2d**, *ρ* = 0.15, *P* = 0.31). Thus, the tRNA/ribosome ratio appears to be tightly constrained in fast-growing bacteria, and is likely regulated through replication-associated gene dosage effects in fast growth.

## Conclusions

We conclude that the tRNA/ribosome ratio appears to be tightly constrained across fast-growing bacteria. Its regulation is likely dominated by replication-associated gene dosage effects in fast growth, implemented through the relative chromosomal positioning of tRNA and ribosomal RNA genes. The objective of this regulation is to not only increase the expression of TCs and ribosomes with growth rate, but to also adjust their relative concentrations according to the RNA composition growth law quantified by equations (1) and (2).

## Methods

### Derivation of the optimal TC/ribosome ratio

In recent work, we have shown that the growth-rate dependent composition of the translation machinery in *E. coli* is accurately described by predictions based on detailed reaction kinetics and the numerical minimization of the total mass of all participating molecules [37]. This minimization was motivated by the observation that the cellular dry mass density is approximately constant across growth conditions [54]; the minimization of the energy consumed or the enzyme mass required for the production of the different molecules led to almost identical results [37].

Here, we consider a much simpler representation of the elongation step of protein translation, which can be modeled as an enzymatic reaction following Michaelis-Menten kinetics [5]. In this case, the minimization of the combined mass concentration of ribosome and TC can be performed analytically, as demonstrated by Dourado *et al.* [9]. At a given reaction flux *v*, the optimal concentration of the ribosome *R* (the “enzyme”) can be written as a function of *v* [9]:

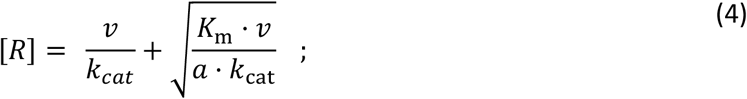

Here, *a* is the ratio of molecular weights of enzyme and substrate, *k*_cat_ is the turnover number of the ribosome, and *K*_m_ is the ribosome’s Michaelis constant for TC. In *E. coli,* the molecular weight of the ribosome is 2307.0 kDa and the molecular weight of a TC is 69.6 kDa [37], thus *a* = 33.1. For a single TC, *K*_m-singleTC_ = 3 μM [5]; the effective number of TC [5] is 34 (the predicted expressed tRNA in Ref. [37]), and thus *K*_m_ = 34 · *K*_m-singleTC_ = 102 μM. *k*_cat_ =22 s^-1^ is the observed maximal translation rate of a ribosome [5]. The optimal concentration of TCs (the “substrate” *M*) can also be expressed as a function of *v*,

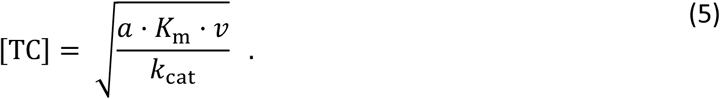

Thus, the TC/ribosome concentration ratio can be written as

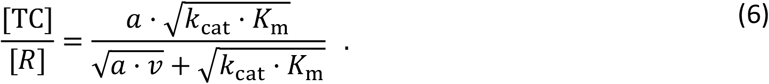

At steady state, protein production rate (*v*) equals to rate of protein dilution by volume growth,

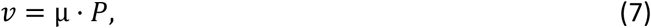

with growth rate *μ* and total cellular protein concentration *P* (in units of amino acids per volume). Thus, equation (6) can be rewritten as (equation (1) of the main text)

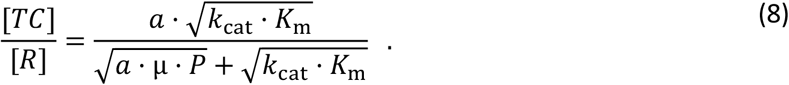

The protein concentration *P* is calculated from *E. coli* proteome expression data [45] and cell volume [55] for growth on glucose,

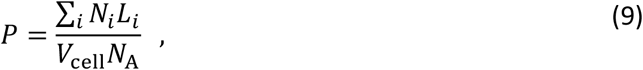

where *N_i_*is the copy number per cell and *L_i_*the length of protein *i* [45], *V*_cell_ is the cell volume [55], and *N*_A_is the Avogadro constant. In a more recent publication [56], the authors of Ref. [55] re-measured the volume of cells by super-resolution microscopy and found that cell volume was overestimated in Ref. [55] (by a factor of 0.67^-1^ for growth on glucose). We thus modified cell volume by a factor of 0.67 relative to the values in Ref. [55], resulting in *P* = 1.16 × 10^6^ μM.

By multiplying the left-hand side of equation (8) with the molecular weight ratio of tRNA to rRNA, we obtain the tRNA and rRNA mass ratio (equation (2) of the main text),

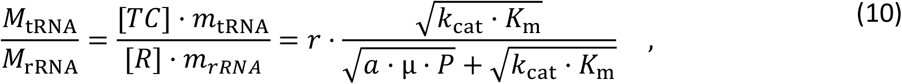

with

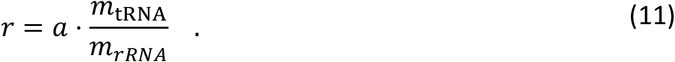

Here, *m*_tRNA_ is the molecular mass of tRNA, *m*_rRNA_ is the total mass of RNA in one ribosome, and *r* is the ratio of the tRNA mass fraction of a TC and the rRNA mass fraction of the ribosome. For bacteria, we use data from *E. coli* (*m*_tRNA_ = 25.8 *kDa, m*_rRNA_ = 1480 k*Da*), resulting in *a* = 33.1 and *r* = 0.58. For eukaryotes, we use data from *S. cerevisiae*, resulting in *a* = 40.3 and *r* = 0.59; the molecular weights of the ribosome (3044.4 kDa), rRNA (1750 kDa), TC (75.6 kDa), and tRNA (25.6 kDa) were calculated from the respective sequences according to the *Saccharomyces* Genome Database [57].

### Gene positions

The chromosomal position of the center of the origin of replication (oriC) for different genomes was obtained from the DoriC database (version 10.0) [58]. The start and end positions of rRNA and tRNA genes were downloaded from the RefSeq database (Release 93, downloaded on April 09, 2019); gene locations were defined as the midpoint between gene start and end. We defined gene position as the relative distance of a gene to oriC, calculated as the shortest distance between the gene and oriC on the circular chromosome, divided by half the length of the chromosome. Gene position ranges from 0 to 1.

For some species, more than one oriC has been annotated [58]. However, we found that all oriCs are very close on the chromosome in these species: the maximal distance between two oriCs is less than 1% of the chromosome length. Thus, different oriCs are expected to have a negligible effect on gene position and we randomly selected one of the oriCs to calculate gene position.

### Maximal growth rate dataset

Minimal doubling times *τ_min_* (in hours) were obtained from Ref. [49] and were converted to maximal growth rates as 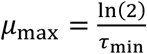. For the analyses, we only used species for which we additionally had genome annotation and oriC location, and which had only one chromosome. The final trimmed dataset contains 170 species (**Additional file 1: Table S3**).

### Gene dosage

We used the Cooper-Helmstetter model [29,30] to calculate gene dosage. The model is briefly summarized below. Let *C* be the time required to replicate the chromosome; let *D* be the time between the termination of a round of replication and the next cell division; let *τ* be the doubling time. The dosage of gene *i* (*X_i_*) is then given by:

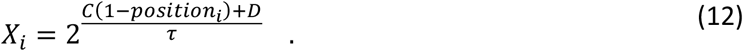

Where *position_i_*is the genomic position of gene *i*. With

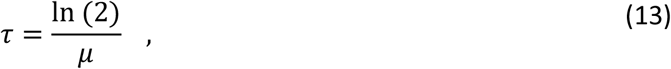

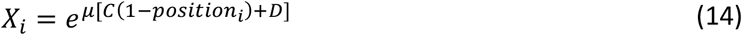

The gene dosage ratio of two genes (*X_i_*/*X_j_*) is then (equation (3) of the main text)

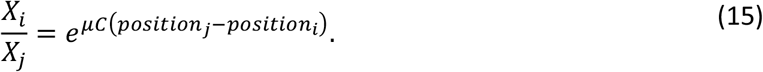

Each genome contains multiple tRNA and rRNA genes. In this case, we use the ratio of the total gene dosages,

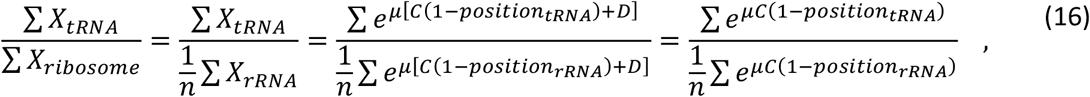

where *n* = 3 is the number of rRNA genes per ribosome.

We assumed a constant DNA replication rate of *k*_rep_ = 1000 bp s^−1^ [27] to calculate the C-period as

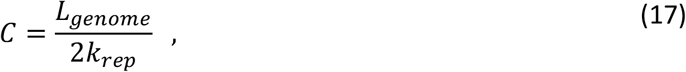

with *L_genome_*the length of the given genome.

## Supplementary information

**Additional file 1: Supplementary Tables**

**Table S1.** Ternary complex per ribosome in *E. coli*.

**Table S2.** tRNA per rRNA in other species.

**Table S3.** The maximal growth rate dataset, including tRNA and rRNA positions, copies, and dosages.

## Declarations

### Ethics approval and consent to participate

Not applicable.

### Consent for publication

Not applicable.

### Availability of data and materials

The experimental data used in this study and the corresponding original publications is provided in Additional file 1.

## Acknowledgments

We thank Hugo Dourado, Peter Schubert, and Deniz Sezer for helpful discussions.

## Author Contributions

XPH designed the study and performed the analyses. XPH and MJL interpreted the results and wrote the manuscript.

